# The brown algae *Saccharina japonica* and *Sargassum horneri* exhibit species-specific responses to synergistic stress of ocean acidification and eutrophication

**DOI:** 10.1101/2021.03.15.435540

**Authors:** Yuxin Liu, Jiazhen Cao, Yaoyao Chu, Yan Liu, Qiaohan Wang, Qingli Gong, Jingyu Li

## Abstract

Ocean acidification and eutrophication are two important environmental stressors. They inevitably impact marine macroalgae, and hence the coastal ecosystem of China. *Saccharina japonica*, as the main culture species in China, is suffering the harmful golden tide caused by *Sargassum horneri*. However, it remains unclear whether the detrimental effects of *S. horneri* on *S. japonica* cultivation become more severe in future acidified and eutrophic scenario. In this study, we respectively investigated the effects of *p*CO_2_ (400 μatm and 1000 μatm) and nutrients (non-enriched and enriched seawater) on the growth, photosynthesis, respiration, chlorophyll contents, and tissue nitrogen of *S. japonica* and *S. horneri*. Results indicated that enrichment of nutrients contributed *S. horneri* to utilize HCO_3_^-^. The carbon acquisition pathway shifted from HCO_3_^-^ to CO_2_ in *S. japonica*, while *S. horneri* remained using HCO_3_^-^ regulated by nutrient enrichment. *S. horneri* exhibited better photosynthetic traits than *S. japonica*, with a higher level of net photosynthetic rate and chlorophyll contents at elevated *p*CO_2_ and enriched nutrients. Tissue nitrogen also accumulated richly in the thalli of *S. horneri* under higher *p*CO_2_ and nutrients. Significant enhancement in growth was only detected in *S. horneri* under synergistic stress. Together, *S. horneri* showed competitive dominance in current study. These findings suggest that increasing risk of golden tide in acidified and eutrophic ocean can most likely result in great damage to *S. japonica* cultivation.

## 1 Introduction

The concentration of atmospheric carbon dioxide (CO_2_) increased approximately 130 pars per million (ppm) since the Industrial Revolution (Joos & Spahni, 2008; AOAN, 2019). Rising atmospheric CO_2_ dissolve in seawater, causing pH reductions and alterations in chemical balances of dissolved inorganic carbon (DIC) (Feely et al., 2004, 2009; Doney et al., 2009). These changes in pH and DIC are ineluctable consequences of rising atmospheric CO_2_, referred to as ocean acidification (OA) (Doney et al., 2009). Anthropogenic CO_2_ emission is rising at the fastest rate after the Industrial Era (Joos & Spahni, 2008; AOAN, 2019), thus leading to a continuing decrease in seawater pH (Feely et al., 2004, 2009; Doney et al., 2009; Feely, Doney & Cooley, 2009). OA significantly affects the physiological processes and ecological functions of seaweeds and other marine organisms (Gazeau et al., 2007; Edmunds, 2011; Koch et al., 2013; Kroeker et al., 2013; Enochs et al., 2015; Gao et al., 2019). Previous studies showed that photosynthetic organisms including macroalgae appear to benefit from elevated CO_2_ and tolerant to declined pH (Shi et al., 2012, 2019; Britton et al., 2016; Gao et al., 2019). A body of evidence indicates that OA actively stimulates the growth of kelps, such as *Saccharina latissima*, *Ulva rigida* and *Macrocystis pyrifera* which were carbon limited in nearshore environment (Swanson & Fox, 2007; Xu et al., 2019; Hurd et al., 2020; Zhang et al., 2020). On the other hand, OA simultaneously reduces the calcification of *Marginopora rossi*, *Porolithon onkodes* and other calcified algae (Reymond et al., 2013; Johnson & Carpenter, 2018).

Furthermore, human pollution, agricultural production and atmospheric deposition have dramatically increased since 1970s, resulting in excessive nutrients input to coastal seawater (Smith et al., 2003; Felipe van der Struijk & Kroeze, 2010; Strokal et al., 2014; Brockmann et al., 2018; Murray et al., 2019). This process leads to another environmental issue known as eutrophication (Smith et al., 2003). Several studies showed that water quality slightly recovered from previous eutrophic state in the Baltic Sea, Chesapeake Bay and other coastal seas (Okino & Kato, 1987; Andersen et al., 2017; McCrackin et al., 2017; Duarte & Krause-Jensen, 2018). In contrast, severe eutrophic areas are still located at some key bays in China, including Liaodong Bay, Yangtze River Estuary and other jurisdictional seas (MEE, 2019). With exceeded nutrients supply, eutrophication can enhance the growth of phytoplankton, fast-growing filamentous and mat-forming opportunistic macroalgae (Pedersen & Borum, 1997; Wernberg et al., 2018). Degraded water quality from eutrophication is critical for the development, persistence and expansion of harmful algae blooms (HABs) (Heisler et al., 2008). Recent reports showed that microalgal blooms, *Ulva*-dominated green tides and *Sargassum*-dominated golden tides have substantially increased worldwide (Glibert et al., 2005; Smetacek & Zingone, 2013; Kudela et al., 2015; Wang et al., 2018). HAB resulted from eutrophication affects substance circulation, primary productivity, community structure and marine ecosystem service (Norkko & Bonsdorff, 1996a,b; Glibert et al., 2005; Rabouille et al., 2008; Heisler et al., 2008; Smetacek & Zingone, 2013; Anderson et al., 2015; Kudela et al., 2015; Watson et al., 2015).

Several studies have found that coral reef systems are negatively affected by OA and nutrient enrichment (Hoegh-Guldberg et al., 2007; Ge et al., 2017; Guan et al., 2020). For phytoplankton, marine diazotrophs such as *Trichodesmium* spp. increase their N2 fixation under elevated CO_2_ in nitrogen enriched cultures (Eichner, Rost & Kranz, 2014; Hutchins & Fu, 2017). However, limited investigations aimed to reveal the ecophysiological effects of OA and eutrophication on marine macrophytes. Previous studies indicated that the growth and quality of *S. japonica* were inhibited and threatened by the interactive effects of OA and eutrophication. (Chu et al., 2019, 2020). In contrast, there was an enhanced production of amino acid and fatty acid in *Ulva* species at elevated CO_2_ concentration and nutrient level (Gao et al., 2018). Thus, the responses to the synergistic stress of OA and eutrophication are species-specific in macroalgae. The rise of acidity in coastal ocean was found to be greater under eutrophication (Cai et al., 2011). This severe scenario potentially aggravate the disappearance of habitat-forming seaweeds worldwide (Wernberg et al., 2018; Filbee-Dexter & Wernberg, 2018). It is thus important to understand how macroalgae will response to the future synergistic stress of OA and eutrophication.

The kelp *Saccharina japonica* is the foremost commercial harvesting alga among northwestern Pacific countries (Kim et al., 2017; Chung, Sondak & Beardall, 2017). In previous studies, the growth, photosynthesis, and nutrient uptake of *S. japonica* were significantly enhanced under elevated CO_2_ concentrations (Swanson & Fox, 2007; Zhang et al., 2020). Also, excess nutrient availability significantly promoted the growth and physiological performance of *S. japonica* (Gao et al., 2017). On the other hand, the sheet-like macroalgae *Sargassum horneri* blooms frequently occur in recent years (Liu et al., 2013; Xiao, 2019), whose floating thalli have caused detrimental impacts on *S. japonica* aquaculture (Xiao, 2019). Many investigations have focused on how environmental factors affect population dynamics and distributions of *S. horneri* in East China Sea and Yellow Sea (Xiao, 2019; Xiao et al., 2020; Choi et al., 2020). However, it remains unclear whether *S. horneri* is more resilient to the synergistic stress of OA and eutrophication than *S. japonica*.

In the present study, we investigated the synergistic stress of OA and eutrophication on growth, photosynthesis, respiration, chlorophyll contents, and tissue nitrogen of sporophytes of *S. japonica* and *S. horneri* appearing in the same period. The results are expected to reveal the species-specific ecophysiological responses of *S. japonica* and *S. horneri*, and determine which alga has greater resilience and interspecific competitive dominance under synergistic stress of OA and eutrophication.

## 2 Materials and Methods

### 2.1 Algal collection and maintenance

The sporophytes of *S. japonica* (approximately 80 cm in average length, n = 20) and *S. horneri* (approximately 150 cm in average length, n = 20) were collected in Rongcheng, Shandong, China (36°07’N, 120°19’E), in December 2019. The *S. japonica* samples were from cultivated populations, with *S. horneri* twining on, or floating between their rafts. The samples were kept in cold foam boxes filled with seawater and quickly transported to the laboratory within 8 h. Healthy sporophytes were selected and rinsed several times with sterilized seawater to remove the epiphytes and detritus. More than 100 discs (1.4 cm in diameter) were punched from the meristem of *S. japonica* with a cork borer, and more than 100 segments (4-5 cm in length) were cut from the apical part of *S. horneri* branches for the subsequent experiments. The discs and segments were maintained separately in plastic tanks containing 3 L filtered seawater. The seawater was renewed daily during the maintenance. These samples were maintained at an irradiance of 90 μmol photons·m^−2^·s^−1^ with a 12L: 12D photoperiod, and 10 □, the seawater temperature of the collection area, for 3 days to reduce the negative effects of excision.

### 2.2 Culture experiment

The culture experiment was conducted over a period of 6 days under combinations two *p*CO_2_ levels (400 μatm and 1000 μatm) and two nutrient levels (non-enriched natural seawater and nutrient-enriched seawater). The nutrient-enriched level was enriched 50% PESI medium (Tatewaki, 1966), which was made by sterilized seawater from coastal Qingdao. There was a total of 4 experimental treatments and each had 3 replicates. Four individuals were cultured in each of 12 gently aerated side-arm flasks, in which each contained 500 mL non-enriched or enriched seawater at 10 □. The culture medium was renewed on the third day of the experiment.

### 2.3 Carbonate chemistry parameters

For the treatments under two *p*CO_2_ levels, the samples were cultured in two CO_2_ incubators (GXZ-380C-C02, Jiangnan Instruments Factory, Ningbo, China). The 400 μatm was achieved by bubbling ambient air. And the 1000 μatm was obtained through gas cylinders of the incubator. The pH value of the medium in each flask was measured by a pH meter (Orion STAR A211; Thermo Scientific). The salinity was measured by a seawater salimeter (0-100‰, Aipli). Other indirectly measured carbonate chemistry parameters of all treatments were calculated based on the pH values, salinity, *p*CO_2_ levels, the equilibrium constants *K*_1_ and *K*_2_ for carbonic acid dissociation, and *K*_B_ for boric acid, using CO2SYS software (Robbins & Kleypas, 2018).

### 2.4 Measurement of growth

The growth of *S. japonica* and *S. horneri* was determined by weighing fresh weight (FW) of discs or thalli. The discs and thalli were gently scrubbed with tissue paper to remove water from the surface before being weighed. The relative growth rate (RGR) was calculated as the following formula:

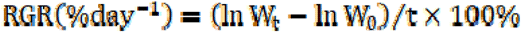

where t is the time period of culture experiment **W_0_**, is the initial FW, **W_t_** is the FW after t days of culture.

### 2.5 Measurement of photosynthesis and respiration

The net photosynthetic rate (P_n_) and the respiration rate (R_d_) of the samples was measured by a manual oxygen meter (FireSting O_2_ □, Pyro Science). After measuring the FW, four discs or segments of each replicates were transferred to the oxygen electrode cuvette with 330 mL culture medium from their own flasks. The medium was magnetically stirred during the measurement to ensure the even diffusion of oxygen. The irradiance and temperature conditions were set the same as the growth chambers. The samples were set to acclimate to the conditions in the cuvette for 5 min before the measurements. The oxygen concentration in the medium was recorded per minute for 10 min. The increase of oxygen content in the medium within 5 min was defined as the P_n_, and the decrease of oxygen content in darkness in the following 5 min was defined as R_d_. The P_n_ and R_d_ were presented as μmol O_2_·min^−1^·g^−1^ FW.

### 2.6 Measurement of chlorophyll contents

Approximately 0.2 g (FW) of the samples from every replicate were used for the extraction of photosynthetic pigments. The discs or segments were dipped in 2 mL dimethyl sulfoxide for 5 min, and the absorbance of supernatant was determined at 665, 631, 582, and 480 nm in the ultraviolet absorbance spectrophotometer (U-2900, HITACHI, Tokyo, Japan). Then the same samples were added 3 mL acetone, setting for 2 h. Before the measurements, 1 mL methanol and 1 mL distilled water was added to the supernatant. The absorbance was obtained at 664, 631, 581, and 470 nm. The contents of chlorophyll (Chl) *a* and *c* were calculated according to the following equation:

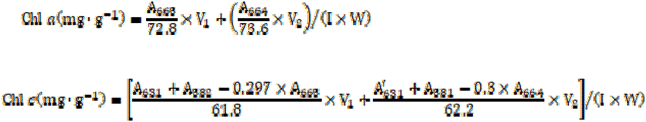

where **A_665_**, **A_664_**, **A_582_** and **A_581_** represent the absorbance at 665, 664, 582 and 581 nm, **A_631_** is the first absorbance at 631 nm, 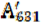 is the second absorbance at 631 nm, **V_1_** is the volume of the first extract, **V_2_** is the volume of the second extract, I is the optical path length, and W is the FW of measured samples.

### 2.7 Measurement of tissue nitrogen

One disc or segment was randomly selected from every replicate for the measurement of tissue nitrogen (TN) contents. The samples were completely dried at 80 □, and ground into powder. About 2-3 mg powder was used to measure the TN contents in the elemental analyzer (Vario EL □, Elementar, Germany). The TN contents were normalized to %DW.

### 2.8 Data analysis

Results were expressed as mean ± standard deviation (n = 3). Prior to the analysis, the data were conformed to a normal distribution (Shapiro-Wilk test, *P* >0.05) and homogeneity of variance (Levene’s test, *P* >0.05). Two-way analysis of variance (ANOVA) was conducted to assess the combined effects of *p*CO_2_ and nutrient levels on carbonate chemistry parameters, RGR, P_n_, R_d_, Chl *a*, Chl *c*, and TN. Tukey honest significance difference (HSD) was conducted to determine the significance levels of factors (*P* <0.05). Pearson correlation coefficient (PCCs) was conducted to analyze the correlations of each experimental indicator with *p*CO_2_ and nutrients levels (*P* <0.05). Data were analyzed in SPSS 22.0 software.

## 3 Results

### 3.1 Carbonate chemistry parameters of culture medium

At the same *p*CO_2_ level, two-way ANOVA showed that nutrients had no significant effects on any parameter (Table 1). In the culture medium of *S. japonica*, elevated *p*CO_2_ decreased the pH by 0.3 and CO_3_^2-^ by 57%, but it increased the DIC by 12%, HCO_3_^-^ by 22%, and CO_2_ by 187% in both the non-enriched and enriched nutrient treatments. In the culture medium of *S. horneri*, elevated *p*CO_2_ decreased the pH by 0.4 in both nutrient levels and CO_3_^2-^ by 75% (non-enriched) and 65% (enriched), but it increased the DIC by 27% (non-enriched) and 4% (enriched), HCO_3_^-^ by 13% (non-enriched) and 5% (enriched), and CO_2_ by 191% in both nutrient treatments.

**Table 1.**
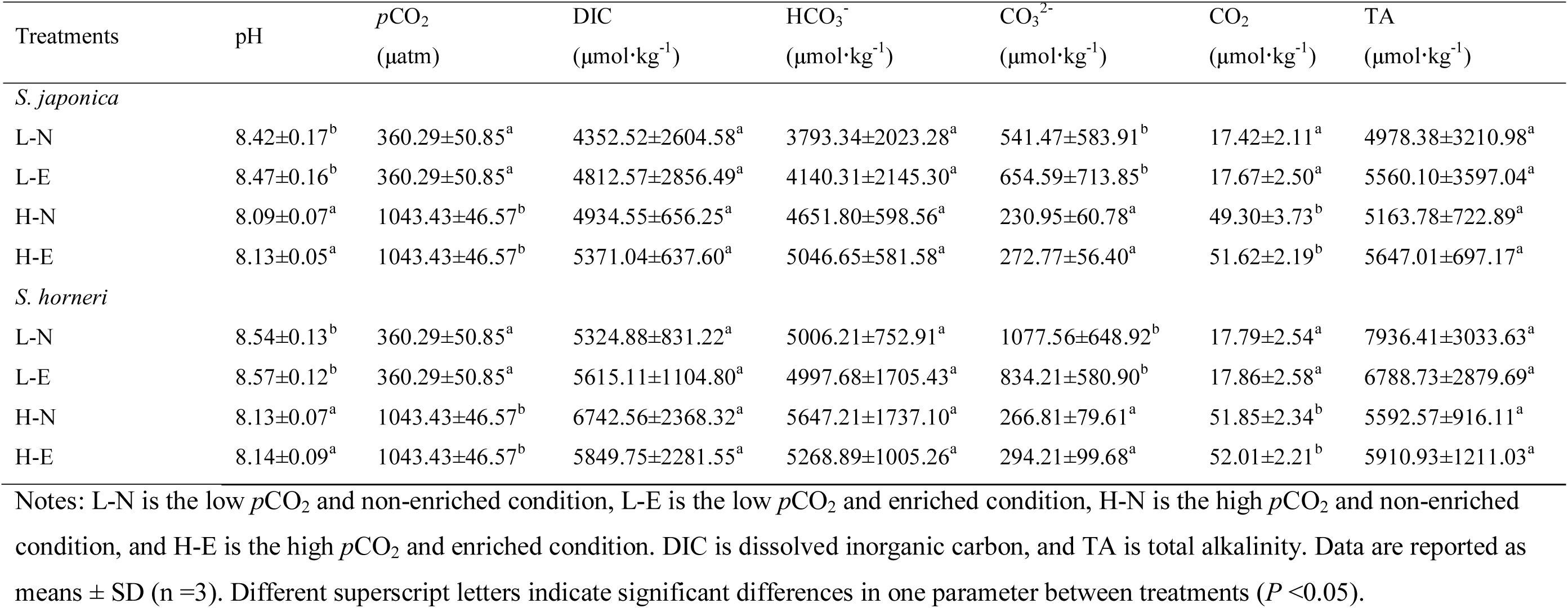
Parameters of the seawater carbonate system at different *p*CO_2_ and nutrient conditions.

**Table 2.** Analysis of variance (two-way ANOVA) examining the statistical differences of experimental parameters of *S. japonica* among *p*CO_2_ and nutrients

| Factors | *S. japonica* |  |  |
| --- | --- | --- | --- |
|  | df | F | P |
| **RGR** |  |  |  |
| $p\text{CO}_2$ | 1 | 0.188 | 0.676 |
| Nutrients | 1 | 0.358 | 0.566 |
| $p\text{CO}_2 \times \text{Nutrients}$ | 1 | 0.115 | 0.743 |
| **$P_n$** |  |  |  |
| $p\text{CO}_2$ | 1 | 0.364 | 0.563 |
| Nutrients | 1 | 5.885 | < 0.05 |
| $p\text{CO}_2 \times \text{Nutrients}$ | 1 | 0.001 | 0.972 |
| **$R_d$** |  |  |  |
| $p\text{CO}_2$ | 1 | 1.739 | 0.224 |
| Nutrients | 1 | 0.039 | 0.848 |
| $p\text{CO}_2 \times \text{Nutrients}$ | 1 | 0.450 | 0.521 |
| **Chl *a*** |  |  |  |
| $p\text{CO}_2$ | 1 | 13.786 | < 0.01 |
| Nutrients | 1 | 9.476 | < 0.01 |
| $p\text{CO}_2 \times \text{Nutrients}$ | 1 | 8.251 | < 0.05 |
| **Chl *c*** |  |  |  |
| $p\text{CO}_2$ | 1 | 0.054 | 0.822 |
| Nutrients | 1 | 9.308 | < 0.05 |
| $p\text{CO}_2 \times \text{Nutrients}$ | 1 | 0.088 | 0.774 |
| **TN** |  |  |  |
| $p\text{CO}_2$ | 1 | 0.042 | 0.843 |
| Nutrients | 1 | 158.903 | < 0.001 |
| $p\text{CO}_2 \times \text{Nutrients}$ | 1 | 0.659 | 0.440 |

**Table 3.** Analysis of variance (two-way ANOVA) examining the statistical differences of experimental parameters of *S. horneri* among *p*CO_2_ and nutrients

| Factors | *S. horneri* |  |  |
| --- | --- | --- | --- |
|  | df | F | P |
| **RGR** |  |  |  |
| $p\text{CO}_2$ | 1 | 0.569 | 0.475 |
| Nutrients | 1 | 4.550 | < 0.05 |
| $p\text{CO}_2 \times \text{Nutrients}$ | 1 | 0.749 | 0.415 |
| **$P_n$** |  |  |  |
| $p\text{CO}_2$ | 1 | 0.025 | 0.879 |
| Nutrients | 1 | 0.286 | 0.607 |
| $p\text{CO}_2 \times \text{Nutrients}$ | 1 | 0.279 | 0.612 |
| **$R_d$** |  |  |  |
| $p\text{CO}_2$ | 1 | 0.015 | 0.904 |
| Nutrients | 1 | 2.607 | 0.145 |
| $p\text{CO}_2 \times \text{Nutrients}$ | 1 | 0.988 | 0.349 |
| **Chl *a*** |  |  |  |
| $p\text{CO}_2$ | 1 | 1.420 | 0.268 |
| Nutrients | 1 | 13.909 | < 0.01 |
| $p\text{CO}_2 \times \text{Nutrients}$ | 1 | 0.077 | 0.789 |
| **Chl *c*** |  |  |  |
| $p\text{CO}_2$ | 1 | 0.592 | 0.463 |
| Nutrients | 1 | 0.740 | 0.414 |
| $p\text{CO}_2 \times \text{Nutrients}$ | 1 | 0.060 | 0.812 |
| **TN** |  |  |  |
| $p\text{CO}_2$ | 1 | 6.053 | < 0.05 |
| Nutrients | 1 | 27.868 | < 0.01 |
| $p\text{CO}_2 \times \text{Nutrients}$ | 1 | 2.147 | 0.181 |

### 3.2 Growth

The differences in *p*CO_2_ and nutrients yielded no significant effects on RGR of *S. japonica*, but nutrients significantly promoted the growth of *S. horneri* (Fig. 1). At both 400 μatm and 1000 μatm, the RGR of *S. japonica* decreased due to enriched nutrient. In contrast, the RGR of *S. horneri* significantly increased in excessive nutrient availability (F = 4.550, *P* < 0.05). PCCs showed that RGR of *S. horneri* positively correlated with both *p*CO_2_ and nutrients. In contrast, RGR of *S. japonica* positively correlated with *p*CO_2_, but negatively correlated with nutrients (Table 4). Together, *S. horneri* showed more promotive growth under the synergistic stress.

**Fig. 1.**
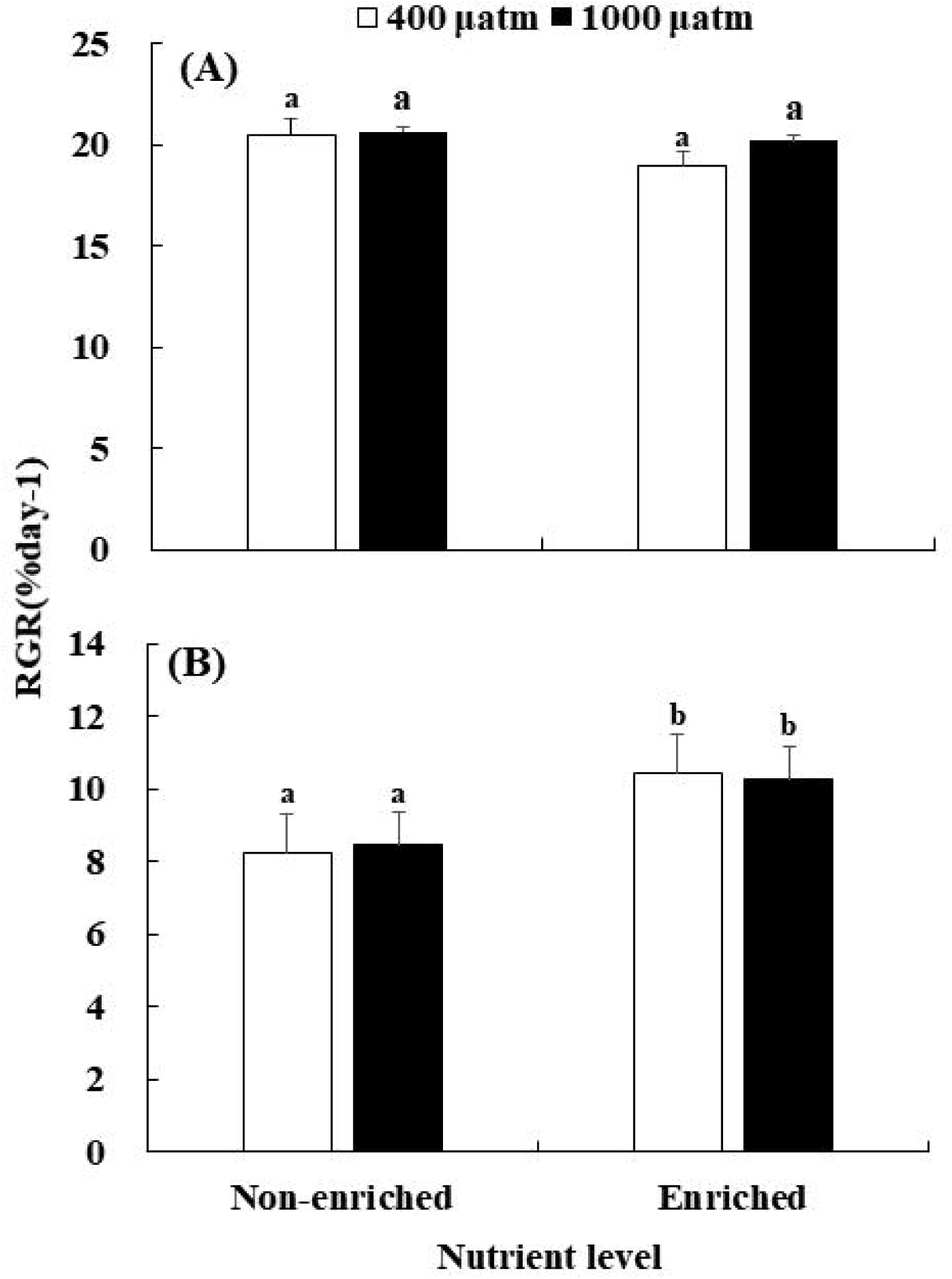
Relative growth rate (RGR) of *S. japonica* (A) and *S. horneri* (B) cultured at different *p*CO_2_ and nutrient conditions for 6 days. Data are reported as means ± SD (n =3). Different letters above the error bars indicate significant differences between treatments (*P* <0.05).

**Table 4.** The Pearson correlation coefficient (PCCs) of various experimental indicators of *S. japonica* and *S. horneri* with *p*CO_2_ and nutrients levels

| Treatments | RGR | P<sub>n</sub> | R<sub>d</sub> | Chl *a* | Chl *c* | TN |
| --- | --- | --- | --- | --- | --- | --- |
| *S. japonica* |  |  |  |  |  |  |
| $p\text{CO}_2$ | 0.533** | 0.241 | 0.883* | 0.359 | 0.380 | -0.016 |
| Nutrients | -0.736** | 0.970** | 0.133* | 0.933* | 0.924* | 0.998** |
| *S. horneri* |  |  |  |  |  |  |
| $p\text{CO}_2$ | 0.021* | 0.030* | 0.670* | 0.463* | 0.953** | 0.410 |
| Nutrients | 0.994** | -0.961** | -0.711** | 0.777** | -0.246** | 0.879* |

Notes: \* indicates significant correlation ( $P < 0.05$ ), \*\* indicates highly significant correlation ( $P < 0.01$ ).

### 3.3 Photosynthesis and respiration

As shown in Fig. 2, nutrient enrichment significantly increased the P_n_ of *S. japonica* at both CO_2_ concentrations (F = 5.885, *P* < 0.05). While no significant effect was detected in *S. horneri*, P_n_ was lower in nutrient-enriched condition. PCCs showed that P_n_ in *S. japonica* had positive correlations with *p*CO_2_ and nutrients. While *S. horneri* positively correlated with *p*CO_2_, but negatively correlated with nutrients (Table 4). Photosynthesis of *S. horneri* was greater than that of *S. japonica* at elevated *p*CO_2_ and nutrients.

**Fig. 2.**
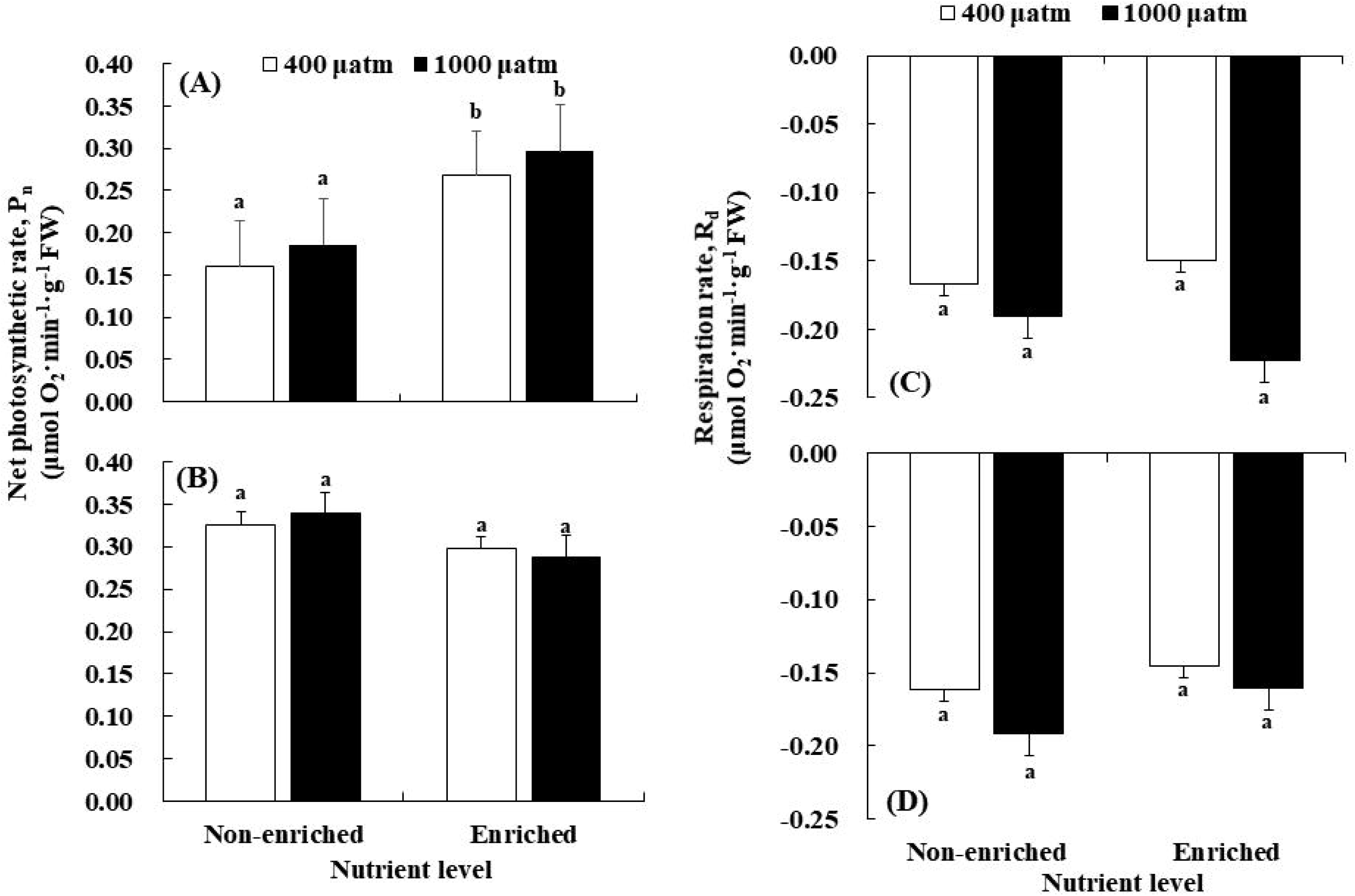
Net photosynthetic rate (P_n_) of *S. japonica* (A) and *S. horneri* (B); Respiration rate (R_d_) of *S. japonica* (C) and *S. horneri* (D) cultured at different *p*CO_2_ and nutrient conditions for 6 days. Data are reported as means ± SD (n =3). Different letters above the error bars indicate significant differences between treatments (*P* <0.05).

The R_d_ in *S. japonica* showed a similar trend to *S. horneri* (Fig. 2). No significant effects on R_d_ of both algae were found in all treatments. At 400 μatm, R_d_ of both species was lower in excess nutrients. The correlation between R_d_ and nutrients of *S. japonica* was positive, but that of *S. horneri* was negative (Table 4). Respiration of *S. japonica* was also greater than that of *S. horneri* under synergistic stress.

### 3.4 Chlorophyll contents

The Chl *a* and *c* contents of *S. japonica* significantly increased under either elevated *p*CO_2_ or enriched nutrient. Both chlorophyll contents reached the maximum under the synergistic stress (Fig. 3). The Chl *a* content of *S. horneri* was significantly increased at enriched nutrients, and also reached the peak in synergistic stress condition. However, the Chl *c* content of *S. horneri* increased only with *p*CO_2_ elevated. Neither *p*CO_2_ nor nutrients significantly affected the Chl *c* in *S. horneri*. PCCs showed positive correlations between Chl *a* with *p*CO_2_ and nutrients in both species. However, correlation between Chl *c* and nutrients was significantly negative in *S. horneri* (Table 4).

**Fig. 3.**
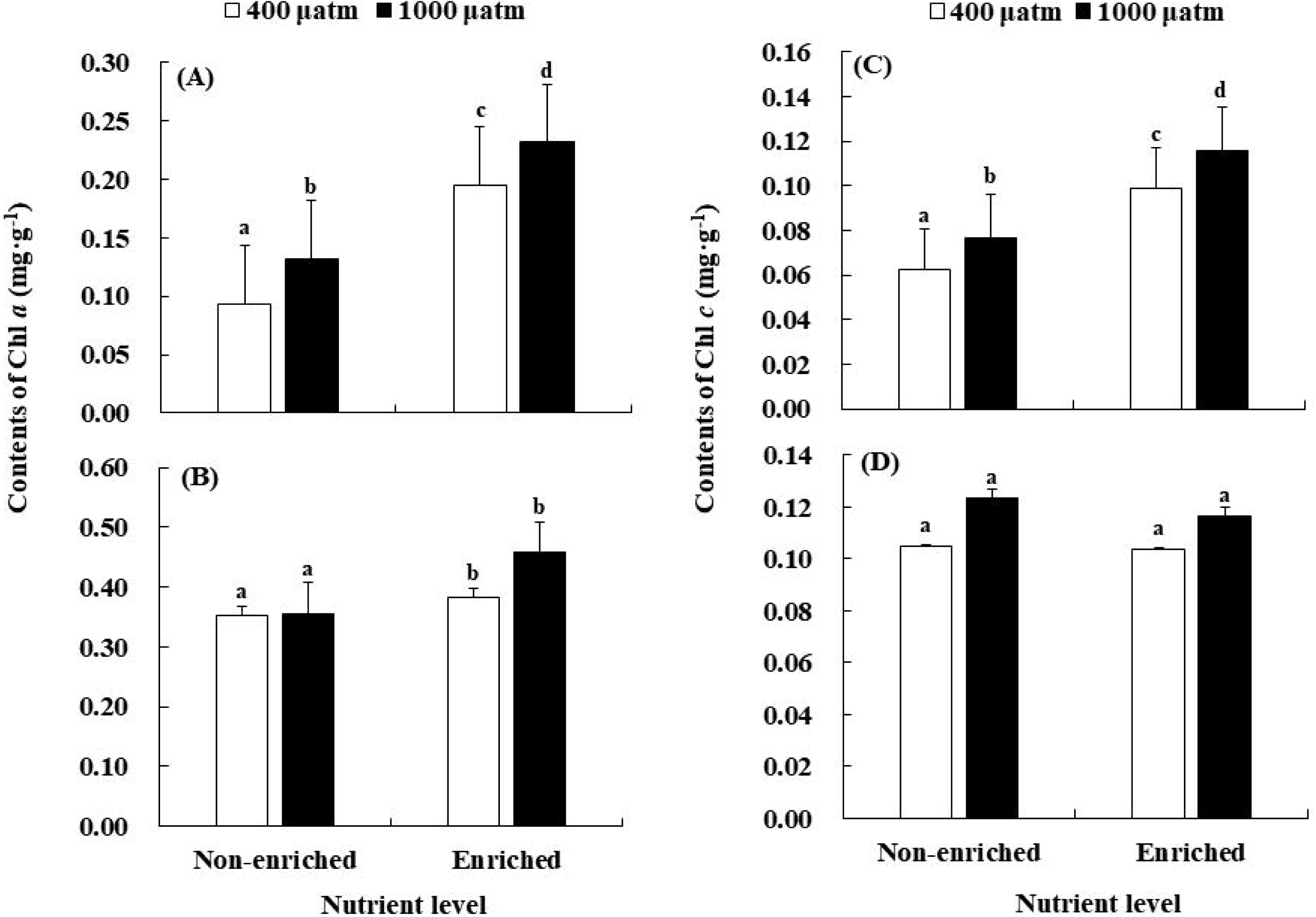
Chl *a* of *S. japonica* (A) and *S. horneri* (B); Chl *c* of *S. japonica* (C) and *S. horneri* (D) cultured at different *p*CO_2_ and nutrient conditions for 6 days. Data are reported as means ± SD (n =3). Different letters above the error bars indicate significant differences between treatments (*P* <0.05).

### 3.5 Tissue nitrogen

The TN contents of *S. japonica* and *S. horneri* significantly increased in nutrient-enriched condition (Fig. 4). Elevated *p*CO_2_ had no significant effect on the TN of *S. japonica*, but significantly promoted the accumulation of TN in *S. horneri*. The correlations between nutrients and TN were significantly positive in the two species. As for the correlations between *p*CO_2_ and TN, it was negative in *S. japonica* but positive in *S. horneri* (Table 4).

**Fig. 4.**
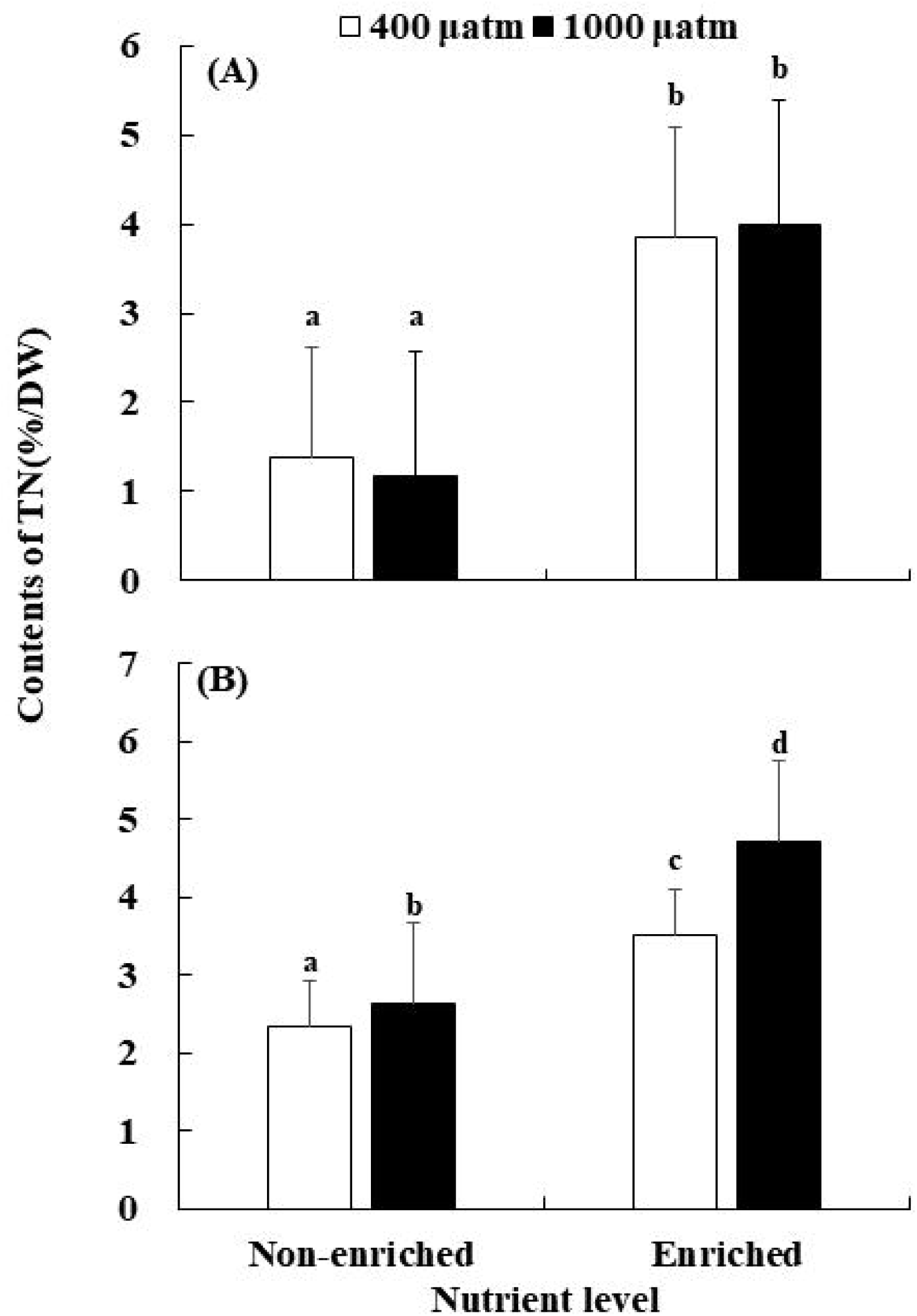
Tissue nitrogen (TN) of *S. japonica* (A) and *S. horneri* (B) cultured at different *p*CO_2_ and nutrient conditions for 6 days. Data are reported as means ± SD (n =3). Different letters above the error bars indicate significant differences between treatments (*P* <0.05).

## 4 Discussion

There was a same increase pattern of DIC in the culture medium of *S. japonica* under two nutrient concentrations, but different case was found in the culture medium of *S. horneri* (Table 1). The effects of the synergistic stress of OA and eutrophication on algae may depend on their precise carbon acquisition pathways utilized. The HCO_3_^-^ in the culture medium of *S. horneri* was lower in enriched nutrient than in non-enriched treatments, indicating more HCO_3_^-^ utilization paralleled with enriched nutrients. Many macroalgae use HCO_3_^-^ rather than dissolved CO_2_ under current seawater *p*CO_2_ concentration (Israel & Hophy, 2002; Badger, 2003; Koch et al., 2013), due to their ribulose-1,5-bisphosphate carboxylase/ oxygenase (Rubisco) is not substrate-saturated at current atmospheric CO_2_ level (Reiskind, Seamon & Bowes, 1988). Marine macroalgae have species-specific responses to elevated CO_2_ because of their various capacities and strategies in CO_2_-concentrating mechanisms (CCMs) to utilize HCO_3_^-^ in seawater (Wu, Zou & Gao, 2008; Raven & Hurd, 2012). Furthermore, DIC acquisition interacts with phosphorus and nitrogen availability (Giordano, Beardall & Raven, 2005), but it remains unclear how *S. horneri* regulates CCMs under excessive nutrient supply. The results indicate that enrichment of nutrients contributed *S. horneri* to the utilization of HCO_3_^-^. When exposed to elevated *p*CO_2_, macroalgae may reduce the use of HCO_3_^-^ by down-regulating their CCMs, and begin to rely on CO_2_ as the primary carbon source (Bjork et al., 1993; Axelsson, Mercado & Figueroa, 2000; Cornwall et al., 2012). This physiological process may have occurred in *S. japonica*, thus leading to the DIC of culture medium remained at the same level after increasing *p*CO_2_ under the two nutrient conditions. In contrast, this study provides an evidence that eutrophication restrains the shift of carbon acquisition pathway in *S. horneri* to cope with higher CO_2_ concentration.

In this study, promotions in RGR were detected in both *S. japonica* and *S. horneri* at elevated *p*CO_2_ although the increases were statistically non-significant. This indicates that both *S. japonica* and *S. horneri* are capable of OA resistance with atmospheric CO_2_ increased to 1000 μatm. To show which algae is competitively dominant under OA condition, we analyzed the P_n_, R_d_, Chl *a*, Chl *c* and TN in both species. The results showed that enhancements to P_n_, R_d_, and chlorophyll contents of *S. japonica* were parallel with *p*CO_2_ elevation. These results are in line with previous investigations on *S. japonica* (Swanson & Fox, 2007; Zhang et al., 2020). The enhancement of P_n_ and chlorophyll contents were also found in other marine macrophytes, including *Gracilariopsis lemaneiformis*, *Pyropia yezoensis* and *Ulva prolifera* (Kang et al., 2017; Li et al., 2018; Bao et al., 2019). However, the P_n_ and chlorophyll contents of *S. horneri* are twice as high as those of *S. japonica*. *S. horneri* increased the utilization of HCO_3_^-^ to maintain its photosynthesis at a higher level. Since P_n_ and Chl *c* of *S. horneri* also increased at 1000 μatm (Fig. 2B, Fig. 3D), photosynthesis of *S. horneri* was further improved on the basis of the original high level. These results indicate that higher photosynthetic level insured *S. horneri* potentially greater resilience to OA in comparison to *S. japonica*.

The significant enhancement in growth was observed in *S. horneri* in nutrient-enriched condition, while no promotion of growth was found in *S. japonica* (Fig.1). In this study, the concentrations of dissolved inorganic nitrate and ammonium were simultaneously increased in nutrient-enriched treatments (Tatewaki, 1966). Increase in nitrogen availability can enhance macroalgae in N uptake rates, tissue N contents, and photosynthetic rates (Valiela et al., 1997). These enhancements accelerate the growth of macroalgae. The significant increase in Chl *a* and TN contents were detected in both species in nutrient-enriched treatments (Fig.3, Fig.4). Previous studies have also determined the same positive physiological responses in *Fucus vesiculosus*, *G. lemaneiformis*, *Hizikia fusiformis* and other macroalgae (Valiela et al., 1997; Kawamitsu & Boyer, 1999; Wu, Zou & Gao, 2008; Raven & Hurd, 2012; Ohlsson et al., 2020). The kinetics of nutrients uptake in macroalgae is affected by the physiological status and the form of nutrients (Raven & Hurd, 2012; Gao et al., 2017). It has been reported that *S. japonica* utilize ammonium first when ammonium and nitrate both exist (Wang et al., 2013), while *S. horneri* firstly takes advantage of nitrate (Yu et al., 2019). We estimated according to the measured ecophysiological traits, because the exact concentrations and formations of nitrogen in culture medium were unclear. *S. horneri* performed higher P_n_, more chlorophyll and TN accumulations under nutrient-enriched condition. Thus, the eutrophic treatment in this study more significantly benefited *S. horneri*, indicating the increased risk of *Sargassum*-dominated golden tide in eutrophic condition.

The current study argued the responses of both *S. japonica* and *S. horneri* under synergistic stress of OA and eutrophication. Significant enhancement in chlorophyll and TN contents was observed in both species (Fig. 3, Fig.4). These results indicated that both *S. japonica* and *S. horneri* improved carbon and nitrogen assimilation. The exceeding nutrient availability in eutrophic scenario regulates these physiological responses in macroalgae to hence the negative effects resulting from declining pH in OA (Young & Gobler, 2016; Chu et al., 2020). However, significant increase in growth was only observed on *S. horneri* (Fig. 1). Increased carbon and nitrogen assimilation in *S. horneri* enhanced its growth more than *S. japonica*. These advantages in ecophyisiological traits may allow *S. horneri* remain dominant and cause damage to *S. japonica* cultivation in future acidified and eutrophic ocean. Furthermore, the damage resulting from golden tide to *S. japonica* cultivation is likely to be more severe in reality. *S.horneri* has vesicles in structure, which can keep the thalli floating and increase carbon acquisition (Smetacek & Zingone, 2013; Choi et al., 2020). Floating *S. horneri* wrap the rafts, shading the cultivated *S. japonica* below (Xiao, 2019; Wu et al., 2019). Thus, we suppose that increasing *S. horneri* biomass shaded cultivated *S. japonica* in a more severe environment with lower light intensity and less carbon availability (Xiao, 2019). The *Sargassum*-dominated golden tide may cause greater damage to *S. japonica* cultivation in acidified and eutrophic ocean. In addition, we need meso-scale experiments to estimate the increasing risk of golden tide in *S. japonica* cultivation.

## 5 Conclusions

It is important to estimate the damage to *S. japonica* cultivation by golden tide resulting from *S. horneri* under the synergistic stress of OA and eutrophication. In this study, we determined that nutrient enrichment contributed *S. horneri* to utilize HCO_3_^-^. *S. horneri* exhibited better photosynthetic traits than *S. japonica*, and tissue nitrogen also accumulated more in thalli of *S. horneri* in elevated *p*CO_2_ and nutrient-enriched treatments. Furthermore, increased carbon and nitrogen assimilation enhanced the growth of *S. horneri* in acidified and eutrophic scenario. Together, *S. horneri* may cause greater damage to *S. japonica* cultivation in acidified and eutrophic ocean.

## Acknowledgements

We sincerely thank Zhu Dasheng, from Shandong Lidao Oceanic Technology Company Limited, for his help in providing algal materials for the experiment. This work is funded by Major Scientific and Technological Innovation Project of Shandong Provincial Key Research and Development Program (2019JZZY020708).

## Notes

### Competing Interest Statement

The authors have declared no competing interest.

### Summary of Updates

We had revised all the figures.

## References

Andersen JH, Carstensen J, Conley DJ, Dromph K, Fleming-Lehtinen V, Gustafsson BG, Josefson AB, Norkko A, Villnäs A, Murray C. 2017. Long-term temporal and spatial trends in eutrophication status of the Baltic Sea. Biological Reviews 92:135–149. DOI: 10.1111/brv.12221.

Anderson CR, Moore SK, Tomlinson MC, Silke J, Cusack CK. 2015. Living with Harmful Algal Blooms in a Changing World: Strategies for Modeling and Mitigating Their Effects in Coastal Marine Ecosystems. In: Coastal and Marine Hazards, Risks, and Disasters. Elsevier Inc., 495–561. DOI: 10.1016/B978-0-12-396483-0.00017-0.

AOAN. 2019. Global Monitoring Laboratory - Carbon Cycle Greenhouse Gases. US Department of Commerce, NOAA, Global Monitoring Laboratory.

Axelsson L, Mercado J, Figueroa F. 2000. Utilization of HCO_3_^-^ at high pH by the brown macroalga *Laminaria saccharina*. European Journal of Phycology 35:53–59. DOI: 10.1080/09670260010001735621.

Badger M. 2003. The roles of carbonic anhydrases in photosynthetic CO_2_ concentrating mechanisms. Photosynthesis Research 77:83–94. DOI: 10.1023/A:1025821717773.

Bao M, Wang J, Xu T, Wu H, Li X, Xu J. 2019. Rising CO_2_ levels alter the responses of the red macroalga *Pyropia yezoensis* under light stress. Aquaculture 501:325–330. DOI: 10.1016/j.aquaculture.2018.11.011.

Bjork M, Haglund K, Ramazanov Z, Pedersen M. 1993. Inducible mechanisms for HCO_3_^-^ utilization and repression of photorespiration in protoplasts and thalli of three species of *Ulva* (Chlorophyta). Journal of Phycology 29:166–173. DOI: 10.1111/j.0022-3646.1993.00166.x.

Britton D, Cornwall CE, Revill AT, Hurd CL, Johnson CR. 2016. Ocean acidification reverses the positive effects of seawater pH fluctuations on growth and photosynthesis of the habitat-forming kelp, Ecklonia radiata. Scientific Reports 6. DOI: 10.1038/srep26036.

Brockmann U, Topcu D, Schütt M, Leujak W. 2018. Eutrophication assessment in the transit area German Bight (North Sea) 2006–2014 – Stagnation and limitations. Marine Pollution Bulletin 136:68–78. DOI: 10.1016/j.marpolbul.2018.08.060.

Cai WJ, Hu X, Huang WJ, Murrell MC, Lehrter JC, Lohrenz SE, Chou WC, Zhai W, Hollibaugh JT, Wang Y, Zhao P, Guo X, Gundersen K, Dai M, Gong GC. 2011. Acidification of subsurface coastal waters enhanced by eutrophication. Nature Geoscience 4:766–770. DOI: 10.1038/ngeo1297.

Choi SK, Oh HJ, Yun SH, Lee HJ, Lee K, Han YS, Kim S, Park SR. 2020. Population dynamics of the “golden tides” seaweed, *Sargassum horneri,* on the southwestern coast of Korea: The extent and formation of golden tides. Sustainability (Switzerland) 12. DOI: 10.3390/su12072903.

Chu Y, Liu Y, Li J, Gong Q. 2019. Effects of elevated *p*CO_2_ and nutrient enrichment on the growth, photosynthesis, and biochemical compositions of the brown alga *Saccharina japonica* (Laminariaceae, Phaeophyta). PeerJ 2019:e8040. DOI: 10.7717/peerj.8040.

Chu Y, Liu Y, Li J, Wang Q, Gong Q. 2020. Nutrient Enrichment Regulates the Growth and Physiological Responses of *Saccharina japonica* to Ocean Acidification. Journal of Ocean University of China 19:895–901. DOI: 10.1007/s11802-020-4359-7.

Chung IK, Sondak CFA, Beardall J. 2017. The future of seaweed aquaculture in a rapidly changing world. European Journal of Phycology 52:495–505. DOI: 10.1080/09670262.2017.1359678.

Cornwall CE, Hepburn CD, Pritchard D, Currie KI, Mcgraw CM, Hunter KA, Hurd CL. 2012. Carbon-use strategies in macroalgae: Differential responses to lowered pH and implications for ocean acidification. Journal of Phycology 48:137–144. DOI: 10.1111/j.1529-8817.2011.01085.x.

Doney SC, Fabry VJ, Feely RA, Kleypas JA. 2009. Ocean acidification: The other CO_2_ problem. Annual Review of Marine Science 1:169–192. DOI: 10.1146/annurev.marine.010908.163834.

Duarte CM, Krause-Jensen D. 2018. Intervention Options to Accelerate Ecosystem Recovery From Coastal Eutrophication. Frontiers in Marine Science 5:470. DOI: 10.3389/fmars.2018.00470.

Edmunds PJ. 2011. Zooplanktivory ameliorates the effects of ocean acidification on the reef coral *Porites* spp. Limnology and Oceanography 56:2402–2410. DOI: 10.4319/lo.2011.56.6.2402.

Eichner M, Rost B, Kranz SA. 2014. Diversity of ocean acidification effects on marine N2 fixers. Journal of Experimental Marine Biology and Ecology 457:199–207. DOI: 10.1016/j.jembe.2014.04.015.

Enochs IC, Manzello DP, Donham EM, Kolodziej G, Okano R, Johnston L, Young C, Iguel J, Edwards CB, Fox MD, Valentino L, Johnson S, Benavente D, Clark SJ, Carlton R, Burton T, Eynaud Y, Price NN. 2015. Shift from coral to macroalgae dominance on a volcanically acidified reef. Nature Climate Change 5:1083–1088. DOI: 10.1038/nclimate2758.

Feely R, Doney S, Cooley S. 2009. Ocean Acidification: Present Conditions and Future Changes in a High-CO_2_ World. Oceanography 22:36–47. DOI: 10.5670/oceanog.2009.95.

Feely RA, Orr J, Fabry VJ, Kleypas JA, Sabine CL, Langdon C. 2009. Present and future changes in seawater chemistry due to ocean acidification. In: Geophysical Monograph Series. American Geophysical Union, 175–188. DOI: 10.1029/2005GM000337.

Feely RA, Sabine CL, Lee K, Berelson W, Kleypas J, Fabry VJ, Millero FJ. 2004. Impact of anthropogenic CO_2_ on the CaCO_3_ system in the oceans. Science 305:362–366. DOI: 10.1126/science.1097329.

Felipe van der Struijk L, Kroeze C. 2010. Future trends in nutrient export to the coastal waters of South America: Implications for occurrence of eutrophication. Global Biogeochemical Cycles 24:1–14. DOI: 10.1029/2009GB003572.

Filbee-Dexter K, Wernberg T. 2018. Rise of Turfs: A New Battlefront for Globally Declining Kelp Forests. BioScience 68:64–76. DOI: 10.1093/biosci/bix147.

Gao K, Beardall J, Häder DP, Hall-Spencer JM, Gao G, Hutchins DA. 2019. Effects of ocean acidification on marine photosynthetic organisms under the concurrent influences of warming, UV radiation, and deoxygenation. Frontiers in Marine Science 6:322. DOI: 10.3389/fmars.2019.00322.

Gao G, Clare AS, Chatzidimitriou E, Rose C, Caldwell G. 2018. Effects of ocean warming and acidification, combined with nutrient enrichment, on chemical composition and functional properties of *Ulva rigida*. Food Chemistry 258:71–78. DOI: 10.1016/j.foodchem.2018.03.040.

Gao X, Endo H, Nagaki M, Agatsuma Y. 2017. Interactive effects of nutrient availability and temperature on growth and survival of different size classes of *Saccharina japonica* (Laminariales, Phaeophyceae). Phycologia 56:253–260. DOI: 10.2216/16-91.1.

Gazeau F, Quiblier C, Jansen JM, Gattuso JP, Middelburg JJ, Heip CHR. 2007. Impact of elevated CO_2_ on shellfish calcification. Geophysical Research Letters 34. DOI: 10.1029/2006GL028554.

Ge C, Chai Y, Wang H, Kan M. 2017. Ocean acidification: One potential driver of phosphorus eutrophication. Marine Pollution Bulletin 115:149–153. DOI: 10.1016/j.marpolbul.2016.12.016.

Giordano M, Beardall J, Raven JA. 2005. CO_2_ concentrating mechanisms in algae: Mechanisms, environmental modulation, and evolution. Annual Review of Plant Biology 56:99–131. DOI: 10.1146/annurev.arplant.56.032604.144052.

Glibert P, Anderson D, Gentien P, Granéli E, Sellner K. 2005. The Global, Complex Phenomena of Harmful Algal Blooms. Oceanography 18:136–147. DOI: 10.5670/oceanog.2005.49.

Guan Y, Hohn S, Wild C, Merico A. 2020. Vulnerability of global coral reef habitat suitability to ocean warming, acidification and eutrophication. Global Change Biology 26:5646–5660. DOI: 10.1111/gcb.15293.

Heisler J, Glibert PM, Burkholder JM, Anderson DM, Cochlan W, Dennison WC, Dortch Q, Gobler CJ, Heil CA, Humphries E, Lewitus A, Magnien R, Marshall HG, Sellner K, Stockwell DA, Stoecker DK, Suddleson M. 2008. Eutrophication and harmful algal blooms: A scientific consensus. Harmful Algae 8:3–13. DOI: 10.1016/j.hal.2008.08.006.

Hoegh-Guldberg O, Mumby PJ, Hooten AJ, Steneck RS, Greenfield P, Gomez E, Harvell CD, Sale PF, Edwards AJ, Caldeira K, Knowlton N, Eakin CM, Iglesias-Prieto R, Muthiga N, Bradbury RH, Dubi A, Hatziolos ME. 2007. Coral reefs under rapid climate change and ocean acidification. Science (New York, N.Y.) 318:1737–1742. DOI: 10.1126/science.1152509.

Hurd CL, Beardall J, Comeau S, Cornwall CE, Havenhand JN, Munday PL, Parker LM, Raven JA, McGraw CM. 2020. Ocean acidification as a multiple driver: How interactions between changing seawater carbonate parameters affect marine life. Marine and Freshwater Research 71:263–274. DOI: 10.1071/MF19267.

Hutchins DA, Fu F. 2017. Microorganisms and ocean global change. Nature Microbiology 2. DOI: 10.1038/nmicrobiol.2017.58.

Israel A, Hophy M. 2002. Growth, photosynthetic properties and Rubisco activities and amounts of marine macroalgae grown under current and elevated seawater CO_2_ concentrations. Global Change Biology 8:831–840. DOI: 10.1046/j.1365-2486.2002.00518.x.

Johnson MD, Carpenter RC. 2018. Nitrogen enrichment offsets direct negative effects of ocean acidification on a reef-building crustose coralline alga. Biology Letters 14. DOI: 10.1098/rsbl.2018.0371.

Joos F, Spahni R. 2008. Rates of change in natural and anthropogenic radiative forcing over the past 20,000 years. Proceedings of the National Academy of Sciences of the United States of America 105:1425–1430. DOI: 10.1073/pnas.0707386105.

Kang JW, Kambey C, Shen Z, Yang Y, Chung IK. 2017. The short-term effects of elevated CO_2_ and ammonium concentrations on physiological responses in *Gracilariopsis lemaneiformis* (Rhodophyta). Fisheries and Aquatic Sciences 20. DOI: 10.1186/s41240-017-0063-y.

Kawamitsu Y, Boyer JS. 1999. Photosynthesis and carbon storage between tides in a brown alga, Fucus vesiculosus. Marine Biology 133:361–369. DOI: 10.1007/s002270050475.

Kim JK, Yarish C, Hwang EK, Park M, Kim Y. 2017. Seaweed aquaculture: Cultivation technologies, challenges and its ecosystem services. Algae 32:1–13. DOI: 10.4490/algae.2017.32.3.3.

Koch M, Bowes G, Ross C, Zhang XH. 2013. Climate change and ocean acidification effects on seagrasses and marine macroalgae. Global Change Biology 19:103–132. DOI: 10.1111/j.1365-2486.2012.02791.x.

Kroeker KJ, Kordas RL, Crim R, Hendriks IE, Ramajo L, Singh GS, Duarte CM, Gattuso JP. 2013. Impacts of ocean acidification on marine organisms: Quantifying sensitivities and interaction with warming. Global Change Biology 19:1884–1896. DOI: 10.1111/gcb.12179.

Kudela RM, Bickel A, Carter ML, Howard MDA, Rosenfeld L. 2015. The Monitoring of Harmful Algal Blooms through Ocean Observing: The Development of the California Harmful Algal Bloom Monitoring and Alert Program. In: Coastal Ocean Observing Systems. Elsevier Inc., 58–75. DOI: 10.1016/B978-0-12-802022-7.00005-5.

Li Y, Zhong J, Zheng M, Zhuo P, Xu N. 2018. Photoperiod mediates the effects of elevated CO_2_ on the growth and physiological performance in the green tide alga *Ulva prolifera*. Marine Environmental Research 141:24–29. DOI: 10.1016/j.marenvres.2018.07.015.

Liu D, Keesing JK, He P, Wang Z, Shi Y, Wang Y. 2013. The world’s largest macroalgal bloom in the Yellow Sea, China: Formation and implications. Estuarine, Coastal and Shelf Science 129:2–10. DOI: 10.1016/j.ecss.2013.05.021.

McCrackin ML, Jones HP, Jones PC, Moreno-Mateos D. 2017. Recovery of lakes and coastal marine ecosystems from eutrophication: A global meta-analysis. Limnology and Oceanography 62:507–518. DOI: 10.1002/lno.10441.

MEE. 2019. Bulletin of Marine Ecology and Environment Status of China in 2018. Beijing.

Murray CJ, Müller-Karulis B, Carstensen J, Conley DJ, Gustafsson BG, Andersen JH. 2019. Past, Present and Future Eutrophication Status of the Baltic Sea. Frontiers in Marine Science 6:2. DOI: 10.3389/fmars.2019.00002.

Norkko A, Bonsdorff E. 1996a. Rapid zoobenthic community responses to accumulations of drifting algae. Marine Ecology Progress Series 131:143–157. DOI: 10.3354/meps131143.

Norkko A, Bonsdorff E. 1996b. Population responses of coastal zoobenthos to stress induced by drifting algal mats. Marine Ecology Progress Series 140:141–151. DOI: 10.3354/meps140141.

Ohlsson LO, Karlsson S, Rupar-Gadd K, Albers E, Welander U. 2020. Evaluation of *Laminaria digitata* and *Phragmites australis* for biogas production and nutrient recycling. Biomass and Bioenergy 140:105670. DOI: 10.1016/j.biombioe.2020.105670.

Okino T, Kato K. 1987. Lake Suwa - Eutrophication and its partial recent recovery. GeoJournal 14:373–375. DOI: 10.1007/BF00208212.

Pedersen M, Borum J. 1997. Nutrient control of estuarine macroalgae:growth strategy and the balance between nitrogen requirements and uptake. Marine Ecology Progress Series 161:155–163. DOI: 10.3354/meps161155.

Rabouille C, Conley DJ, Dai MH, Cai WJ, Chen CTA, Lansard B, Green R, Yin K, Harrison PJ, Dagg M, McKee B. 2008. Comparison of hypoxia among four river-dominated ocean margins: The Changjiang (Yangtze), Mississippi, Pearl, and Rhône rivers. Continental Shelf Research 28:1527–1537. DOI: 10.1016/j.csr.2008.01.020.

Raven JA, Hurd CL. 2012. Ecophysiology of photosynthesis in macroalgae. In: Photosynthesis Research. Springer, 105–125. DOI: 10.1007/s11120-012-9768-z.

Reiskind JB, Seamon PT, Bowes G. 1988. Alternative Methods of Photosynthetic Carbon Assimilation in Marine Macroalgae. Plant Physiology 87:686–692. DOI: 10.1104/pp.87.3.686.

Reymond CE, Lloyd A, Kline DI, Dove SG, Pandolfi JM. 2013. Decline in growth of foraminifer *Marginopora rossi* under eutrophication and ocean acidification scenarios. Global Change Biology 19:291–302. DOI: 10.1111/gcb.12035.

Robbins LL, Kleypas J. 2018. CO_2_ system Calculations.

Shi D, Hong H, Su X, Liao L, Chang S, Lin W. 2019. The physiological response of marine diatoms to ocean acidification: differential roles of seawater CO_2_ and pH. Journal of Phycology 55:521–533. DOI: 10.1111/jpy.12855.

Shi D, Kranz SA, Kim JM, Morel FMM. 2012. Ocean acidification slows nitrogen fixation and growth in the dominant diazotroph Trichodesmium under low-iron conditions. Proceedings of the National Academy of Sciences of the United States of America 109:E3094–E3100. DOI: 10.1073/pnas.1216012109.

Smetacek V, Zingone A. 2013. Green and golden seaweed tides on the rise. Nature 504:84–88. DOI: 10.1038/nature12860.

Smith S V., Swaney DP, Talaue-McManus L, Bartley JD, Sandhei PT, McLaughlin CJ, Dupra VC, Crossland CJ, Buddemeier RW, Maxwell BA, Wulff F. 2003. Humans, hydrology, and the distribution of inorganic nutrient loading to the ocean. BioScience 53:235–245. DOI: 10.1641/0006-3568(2003)053[0235:HHATDO]2.0.CO;2.

Strokal M, Yang H, Zhang Y, Kroeze C, Li L, Luan S, Wang H, Yang S, Zhang Y. 2014. Increasing eutrophication in the coastal seas of China from 1970 to 2050. Marine Pollution Bulletin 85:123–140. DOI: 10.1016/j.marpolbul.2014.06.011.

Swanson AK, Fox CH. 2007. Altered kelp (Laminariales) phlorotannins and growth under elevated carbon dioxide and ultraviolet-B treatments can influence associated intertidal food webs. Global Change Biology 13:1696–1709. DOI: 10.1111/j.1365-2486.2007.01384.x.

Tatewaki M. 1966. Formation of a Crustaceous Sporophyte with Unilocular Sporangia in Scytosiphon lomentaria. Phycologia 6:62–66. DOI: 10.2216/i0031-8884-6-1-62.1.

Valiela I, McClelland J, Hauxwell J, Behr PJ, Hersh D, Foreman K. 1997. Macroalgal blooms in shallow estuaries: Controls and ecophysiological and ecosystem consequences. Limnology and Oceanography 42:1105–1118. DOI: 10.4319/lo.1997.42.5_part_2.1105.

Wang B, Xin M, Wei Q, Xie L. 2018. A historical overview of coastal eutrophication in the China Seas. Marine Pollution Bulletin 136:394–400. DOI: 10.1016/j.marpolbul.2018.09.044.

Wang Y, Xu D, Fan X, Zhang X, Ye N, Wang W, Mao Y, Mou S, Cao S. 2013. Variation of photosynthetic performance, nutrient uptake, and elemental composition of different generations and different thallus parts of *Saccharina japonica*. Journal of Applied Phycology 25:631–637. DOI: 10.1007/s10811-012-9897-y.

Watson SB, Whitton BA, Higgins SN, Paerl HW, Brooks BW, Wehr JD. 2015. Harmful Algal Blooms. In: Freshwater Algae of North America: Ecology and Classification. Elsevier Inc., 873–920. DOI: 10.1016/B978-0-12-385876-4.00020-7.

Wernberg T, Krumhansl K, Filbee-Dexter K, Pedersen MF. 2018. Status and trends for the world’s kelp forests. In: World Seas: An Environmental Evaluation Volume III: Ecological Issues and Environmental Impacts. Elsevier, 57–78. DOI: 10.1016/B978-0-12-805052-1.00003-6.

Wu H, Feng J, Li X, Zhao C, Liu Y, Yu J, Xu J. 2019. Effects of increased CO_2_ and temperature on the physiological characteristics of the golden tide blooming macroalgae *Sargassum horneri* in the Yellow Sea, China. Marine Pollution Bulletin 146:639–644. DOI: 10.1016/j.marpolbul.2019.07.025.

Wu HY, Zou DH, Gao KS. 2008. Impacts of increased atmospheric CO_2_ concentration on photosynthesis and growth of micro- and macro-algae. Science in China, Series C: Life Sciences 51:1144–1150. DOI: 10.1007/s11427-008-0142-5.

Xiao J. 2019. Interim report on progress of drifting Sargassum horneri in Yellow Sea (Genetic diversity of benthic and floating populations of Sargassum in western Yellow Sea).

Xiao J, Wang Z, Song H, Fan S, Yuan C, Fu M, Miao X, Zhang X, Su R, Hu C. 2020. An anomalous bi-macroalgal bloom caused by *Ulva* and *Sargassum* seaweeds during spring to summer of 2017 in the western Yellow Sea, China. Harmful Algae 93:101760. DOI: 10.1016/j.hal.2020.101760.

Xu D, Brennan G, Xu L, Zhang XW, Fan X, Han WT, Mock T, McMinn A, Hutchins DA, Ye N. 2019. Ocean acidification increases iodine accumulation in kelp based coastal food webs. Global Change Biology 25:629–639. DOI: 10.1111/gcb.14467.

Young CS, Gobler CJ. 2016. Ocean Acidification Accelerates the Growth of Two Bloom-Forming Macroalgae. PLoS ONE 5:e0155152. DOI: 10.1371/journal.pone.0155152.

Yu J, Li J, Wang Q, Liu Y, Gong Q. 2019. Growth and Resource Accumulation of Drifting *Sargassum horneri* (Fucales, Phaeophyta) in Response to Temperature and Nitrogen Supply. Journal of Ocean University of China 18:1216–1226. DOI: 10.1007/s11802-019-3835-4.

Zhang X, Xu D, Guan Z, Wang S, Zhang Y, Wang W, Zhang X, Fan X, Li F, Ye N. 2020. Elevated CO_2_ concentrations promote growth and photosynthesis of the brown alga *Saccharina japonica*. Journal of Applied Phycology 32:1949–1959. DOI: 10.1007/s10811-020-02108-1.

